# Prodromal pathogenesis of CLN7 Batten Disease revealed by multimodal biomarkers in macaques

**DOI:** 10.64898/2025.12.02.691930

**Authors:** William A Liguore, Winde Hilven, Lili Lurye, Jordy Akkermans, Marion Decrop, Robert Zweig, Larry S. Sherman, Jodi L McBride, Daniele Bertoglio, Alison R Weiss

**Author notes:** Authors contributed equally (William A Liguore, Winde Hilven). Senior authors contributed equally to this work (Daniele Bertoglio, Alison R Weiss). Corresponding authors: Alison R. Weiss, PhD, Division of Neuroscience, ONPRC; Beaverton, OR 97006, USA, Daniele Bertoglio, PhD, Bio-Imaging Lab, University of Antwerp, Universiteitsplein 1, Wilrijk, Belgium.

## Abstract

Neuronal ceroid lipofuscinosis type 7 (CLN7) is a devastating paediatric neurodegenerative disorder with no cure and limited natural history data to guide therapeutic development. Here, we present the first multimodal characterization of prodromal and early-stage CLN7 disease in Japanese macaques carrying a spontaneous *CLN7*^−/−^ mutation.

Using structural T_2_-weighted MRI for volumetry, [^18^F]FDG PET for glucose metabolism, and [^11^C]PBR28 PET for neuroinflammation, we observed region-dependent patterns in volumetric, molecular, and metabolic alterations.

MRI confirmed the presence of disease-associated atrophy in many cortical and subcortical brain regions, consistent with human pathology, [^18^F]FDG PET revealed early cortical and subcortical widespread hypometabolism, and [^11^C]PBR28 PET imaging detected progressive neuroinflammation in the same brain areas. CSF analyses further showed age-dependent increases in neurofilament light (NfL), providing convergent evidence for neurodegeneration.

Together, these results define a prodromal trajectory in CLN7 disease, establish sensitive imaging and fluid biomarkers, and validate the macaque model as a powerful platform for testing interventions.

## Introduction

Neuronal ceroid lipofuscinosis (NCL) is a rare heterogeneous autosomal recessive disease caused by a genetic mutation in one of 14 identified genes.^1–3^ It is characterized by a period of normal childhood development before symptom onset, when neurological deterioration emerges.^4–8^ Disease subtypes (CLN1-14) vary in severity and timing but share many features including accumulation of lipopigment in the CNS and retina, neuronal loss, and neuroinflammation.^9–13^

Children with CLN7 disease^6^ typically do not survive past their teens.^10,14,15^ Due to its rarity, data characterizing the natural history of affected individuals is scant.^16^ Anatomical MRI studies at advanced disease stages in CLN7 patients have identified the cerebellum, thalamus, and cerebral cortex as brain structures most vulnerable to atrophy.^16^ However, interpretation of these data is limited by the sparse number of studies, single imaging modality, and challenges in diagnosing patients during the prodromal stage leading to lack of data covering the entire disease course. To overcome the limitations of these clinical studies, and to provide insight into region-specific patterns, CLN7 animal models are of great utility for better understanding the mechanisms of disease progression and defining multimodal biomarkers, particularly during prodromal stages.

CLN7 knockout mice mimic some pathological features seen in CLN7 patients, including lipofuscin in the basal ganglia and thalamus.^2,11,17^ However, differences in neurodevelopmental trajectories, brain morphology, and motor function between humans and mice somewhat limits the translational utility of murine models.^11^ Furthermore, species differences in brain size and complexity also limit the degree to which gene therapy vector biodistribution and intra-thecal dosing can be translated between rodents and humans.^12,18–22^

Nonhuman primates (NHP) are of high value due to their close phylogenetic proximity to humans,^19^ and similarities in neuronal development, anatomy, and neuronal circuitry^20^; yet the availability of NHP models has historically been limited to CLN2.^12,21,22^ Recently, a population of Japanese macaques (*Macaca fuscata)* at the Oregon National Primate Research Center (ONPRC) were identified with a naturally occurring *CLN7*^−/−^ mutation identical to those reported in human patients^23^. Preliminary work in these CLN7 macaques revealed a characteristic phenotype in which affected animals displayed progressive neurological deficits (visual impairment, tremor, incoordination, ataxia, and impaired balance),^23^ as well as storage‐ mediated retinal degeneration.^24^ Our prior study, using T_2_-weighted (T_2_w) MRI, demonstrated marked cerebellar and whole cerebrum volume loss, ventricular enlargement and subdural CSF accumulation that was particularly pronounced in animals between the ages of 5–6 years, but the in vivo analyses lacked sub-regional specificity.^23^ Postmortem analyses of advanced cases confirmed widespread accumulation of autofluorescent ceroid lipopigment in brain, cerebellum and cardiac tissue, accompanied by neuronal loss, white matter fragmentation and robust astrogliosis and microglial activation.^23^

Although this work validated key features of advanced CLN7 disease pathology, important gaps remain in our understanding of the prodromal period of this disorder in presymptomatic CLN7 macaques. To address these needs, we characterized the spatiotemporal impact of the *CLN7*^−/−^ mutation in macaques on neurodegeneration and neuroinflammation during earlier stages. To accomplish this, we (i) charted region-specific volumetric alterations; (ii) evaluated cerebral glucose metabolism using [^18^F]FDG PET; (iii) quantified neuroinflammation using [^11^C]PBR28 PET; and (iv) evaluated the utility of several CSF biomarkers of neurodegeneration. Our findings reveal unprecedented insight into region-specific patterns of molecular and structural alterations in CLN7 disease, provide a basis for understanding prodromal CLN7 pathophysiology, and identify potential biomarkers that could be used for evaluating the efficacy of interventions during clinical trials.

## Materials and Methods

### Overview of model and experimental design

This work aimed to perform the first multimodal characterization of CLN7 disease in Japanese macaques carrying a spontaneous *CLN7*^−/−^ mutation. A total of 22 Japanese macaques, (9 females, F;13 males, M) age range 1.4-5.8 years, were included in the study. Seven of these animals carried the spontaneous *CLN7*^−/−^ mutation (2F, 5M) and 15 of these animals were wild type (WT) controls (7F, 8M). No CLN7 heterozygotes (*CLN7*^*+/-*^) were included. As has been described previously,^23^ CLN7 animals typically do not develop clinical signs before they are approximately 3.5-4 years of age. Between 4 and 5 years, mild tremors, altered gait/balance, and visual deficits may be detected when animals are closely observed. Symptom severity increases gradually, so animals often adapt and continue to thrive in living conditions that are well-lit and easy to navigate. At ~5 years, signs of cerebellar ataxia and limited vision are obvious, but manageable. In some cases, animals may also develop seizures or other abnormal behaviours. By ~5.5-6 years, neurological symptoms are severe enough where a humane endpoint is reached.

All animals involved in this study were housed either in outdoor enclosures in social groups, or in indoor enclosures with a socially compatible conspecific. Indoor-housed animals were maintained on a 12-h on/12-h off lighting schedule. Food rations (Purina 5000 Monkey Chow and fresh produce) were provided twice daily, and water was provided *ad libitum*. Animal welfare was in accordance with all United States federal regulations and guidelines established by the National Institutes of Health and the Institutional Animal Care and Use Committee at the ONPRC.

Detailed descriptions of all methods are provided in Supplementary materials. Briefly, to investigate regional brain volumetric alterations, we collected 3D T_2_w imaging sequences using a Siemens Prisma whole-body 3 Tesla (T) MRI instrument (Erlangen, Germany) with a 16-channel paediatric head RF coil. We evaluated cerebral glucose metabolism and quantified neuroinflammation via [^18^F]FDG and [^11^C]PBR28 PET imaging respectively, using a GE Discovery MI 710 PET-CT imaging system. Additionally, in a subset of animals (CLN7, *n*=6; WT, *n*=8), 1 mL of CSF was collected under isoflurane anaesthesia by cisterna magna aspiration and used to measure neurofilament light chain (NfL), glial fibrillary acidic protein (GFAP), total Tau, and ubiquitin carboxy-terminal hydrolase L1 (UCHL1) using ELISA (Simoa platform). The spontaneous nature of this model made it challenging to set up a balanced longitudinal study for all included subjects. As a result, we collected opportunistic samples from animals over time. A comprehensive overview of each modality and time points every animal underwent are provided in Supplementary Tables 1-4. All metrics obtained were investigated at the group level to compare between genotypes, and at the single level to explore the relationship between a given metric with age, reflecting disease progression. Investigators performing all analyses were blinded to genotype until completion of study. Randomization was applied to each experiment. No samples correctly acquired were excluded from the study. Detailed descriptions of the statistical analyses are provided in Supplementary materials.

## Results

### Brain volumes progressively decrease with Batten Disease pathology

To investigate brain atrophy associated with the *CLN7*^−/−^ mutation, MRI-based volumetric analyses were performed across three developmental age bins that captured key stages of disease progression: <2.5 years (younger/pre-symptomatic), 2.5-3.5 years (incipient disease), >3.5 years (clinical stage). Representative T2w images across these stages are shown in Fig. 1A.

**Figure 1.**
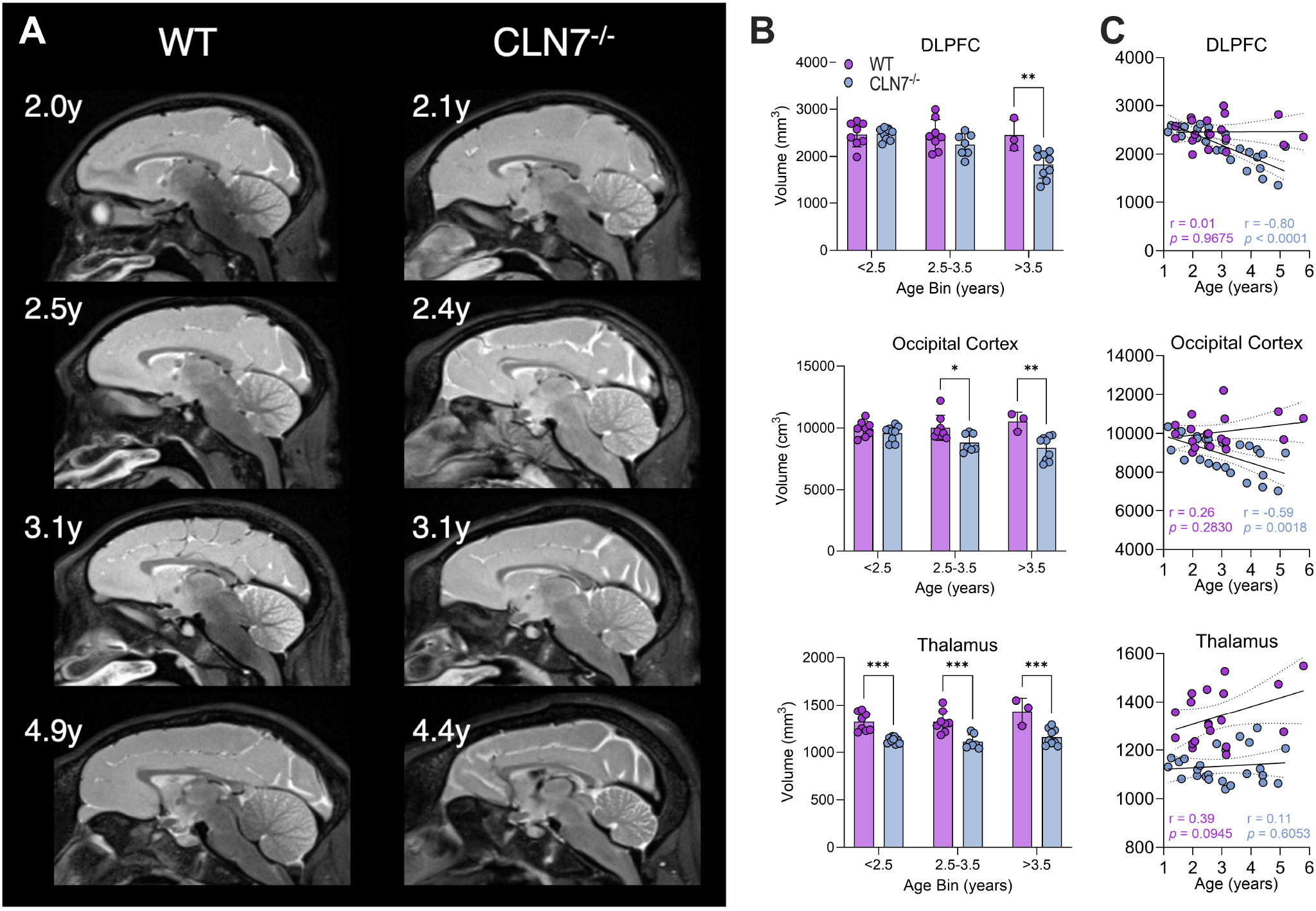
MRI brain volume measurements detected reduced volumes in BD animals only after 4 years of age. **(A)** Panel of sagittal T_2_w MRI brain images of WT Japanese macaques and CLN7 animals at different ages. **(B)** Volumetric analysis comparing WT and CLN7 animals at three age bins. \**P*<0.05, \*\**P*<0.01, \*\*\**P*<0.001. **(C)** Scatterplots displaying volumes over age with separate linear regressions for CLN7 and WT animals. A solid line of best fit is plotted along with 95% confidence intervals in dashed lines. Abbreviations: DLPFC = Dorsolateral Prefrontal Cortex, OCC = Occipital Cortex

To quantify the impact of age and genotype on regional brain structure, two-way ANOVAs were conducted separately for 27 regions of interest (ROIs) in the ONPRC18 gray matter labelmap.^25^ Significant age x genotype interactions were observed in 18 of 27 ROIs (all *P*<0.05), indicating differential age-related trajectories between CLN7 and WT animals. Main effects of genotype were present in 19 ROIs, and main effects of age in 10 ROIs (all *P***<**0.05, see **Supplementary Table 5** for statistical details). Post hoc comparisons between CLN7 and WT animals revealed different patterns across age bins: At the younger/pre-symptomatic stage (<2.5y), significant genotype differences were detected in 3 of 27 ROIs, during the incipient state (2.5-3.5y) in 5 of 27s ROIs, and by the clinical stage (>3.5y) in 22 of 27 ROIs (all *P*<0.05, see **Supplementary Table 5** for statistical details). These findings indicate that regional differences become increasingly widespread as disease advances.

Consistent with this trajectory, Pearson correlation analyses revealed significant negative associations between ROI volume and age in 20 of 27 ROIs in CLN7 animals. In contrast, WT animals exhibited only two significant positive associations between ROI volume and age (all *P*<0.05, see **Supplementary Table 6** for statistical details). The most pronounced age-related volume declines in CLN7 animals occurred in prefrontal, cingulate, sensorimotor, thalamic, and cerebellar regions, consistent with the selective vulnerability of these structures in human CLN7 disease.

### *CLN7*^*-/-*^ mutation leads to widespread alterations in cerebral glucose metabolism

Given the emergence of neurological phenotype around 3.5-4y, we investigated alterations in glucose metabolism due to the *CLN7*^−/−^ mutation using [^18^F]FDG PET imaging. In PET scans collected before the age of 2.5y (Fig. 2A), statistical analysis revealed main effects of genotype (F_(1,40)_=44.35, *P*<0.0001) and brain regions (F_(7,40)_=6.268, *P*<0.0001). Post-hoc group testing identified significantly altered SUVglc values in CLN7 animals compared to WT controls in Frontal Cortex (*P*=0.0035) and Motor Cortex (*P*=0.0376) (Fig. 2B).

**Figure 2.**
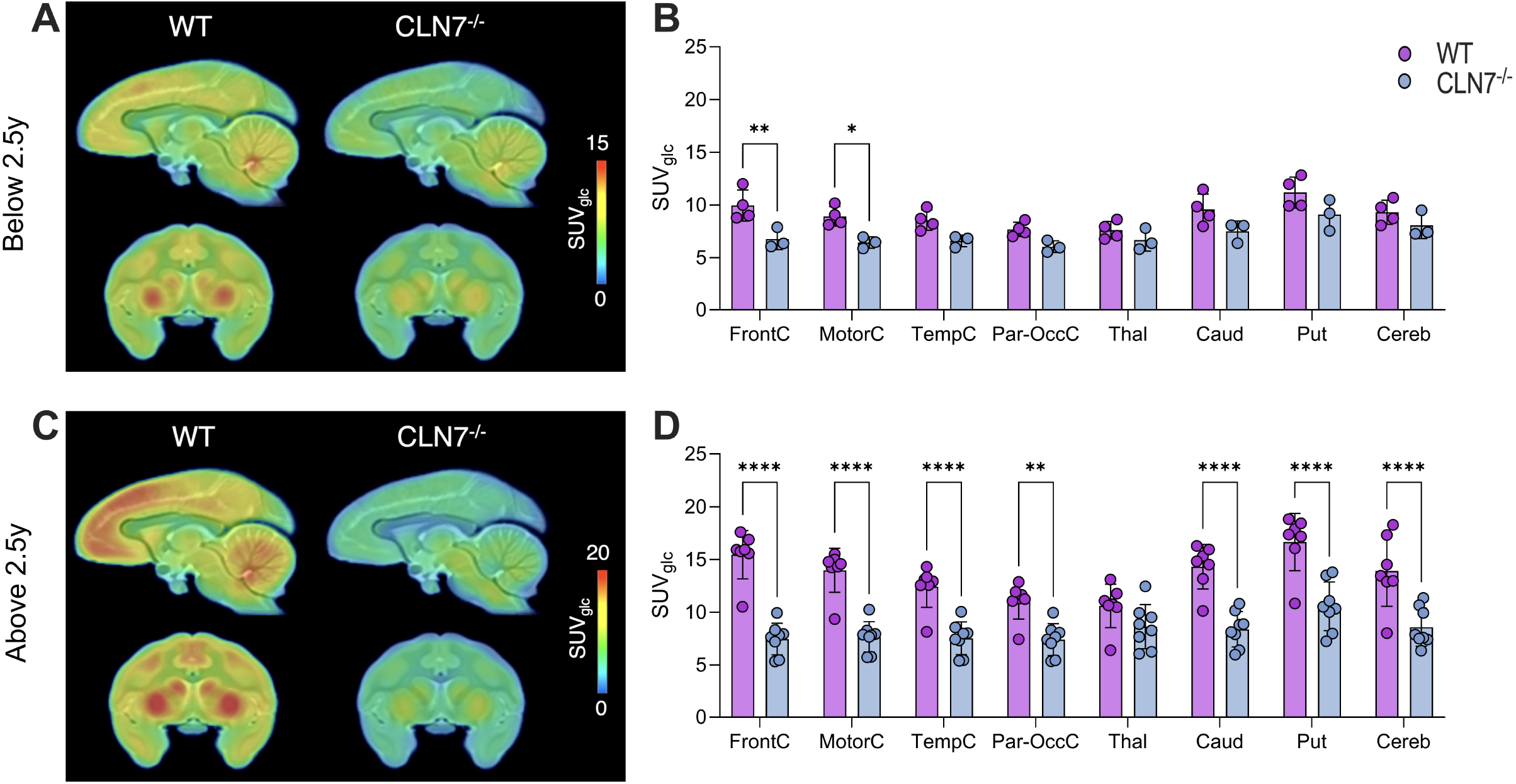
[^18^F]FDG PET imaging revealed reduced cerebral glucose metabolism in CLN7 animals, especially after 2.5 years of age. **(A)** [^18^F]FDG SUV_glc_ images of WT and CLN7^−/−^ animals <2.5 years of age overlaid on ONPRC18 T_2_w MRI template. **(B)** Histogram comparing the SUV_glc_ of WT and CLN7 animals ≤2.5y. **(C)** [^18^F]FDG SUV_glc_ images of WT and CLN7^−/−^ animals >2.5y overlaid on ONPRC18 T_2_w MRI template. **(D)** Histogram comparing SUV_glc_ of WT and CLN7 animals >2.5 y. \**P*<0.05, \*\**P*<0.01, \*\*\*\**P*<0.0001. Abbreviations: FrontC = Frontal Cortex, MotorC = Motor Cortex, TempC = Temporal Cortex, Par-OccC = Parieto-Occipital Cortex, Thal = Thalamus, Caud = Caudate, Put = Putamen, Cereb = Cerebellum.

Additionally, PET scans collected 2.5 years of age and older (Fig. 2C) showed significant main effects of genotype (F_(1,112)_=198, *P*<0.0001), brain region (F_(7,112)_=6.714, *P*<0.0001) as well as interaction (F_(7,112)_=2.956, *P*=0.0072). Post-hoc group analysis indicated a wide-spread statistically significant difference between CLN7 and control in all investigated brain regions except for Thalamus (Fig. 2D).

Finally, we explored the relationship between altered glucose metabolism and age. Correlation analyses indicated a significant positive association in all cortical and sub-cortical regions except Cerebellum for WT animals (all *P*<0.05), whereas this significant increase in glucose metabolism with neurodevelopment was absent in the CLN7 group (all *P*>0.05). (Supplementary Fig. 6).

### Batten Disease pathology results in progressive neuroinflammation

Next, we assessed the presence of neuroinflammation in this CLN7 model using [^11^C]PBR28 PET imaging (Fig. 3A). Statistical analysis revealed a significant main effect of genotype (F_(1,104)_=35.20, *P*<0.0001) with no effect of brain region or interaction. Post-hoc group analysis indicated a statistically significant difference between groups in Thalamus (*P*=0.0137) and Cerebellum (*P*=0.0025) (Fig. 3B). Noteworthy, when correlating SUV with age to account for the large spread of ages in the BD population, significant (*P*<0.05) positive associations with age were found in all regions except the thalamus (Fig. 3C).

**Figure 3.**
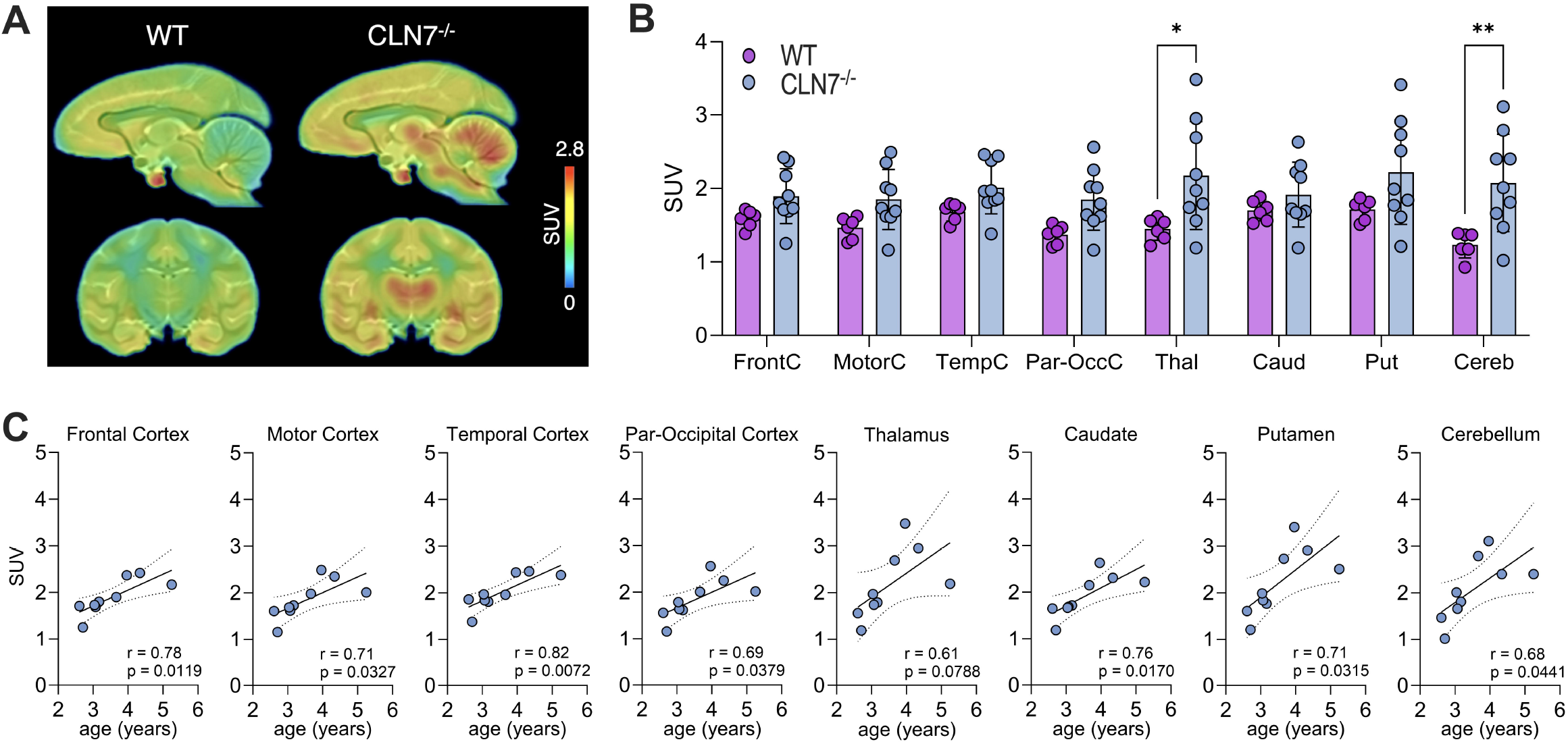
[^11^C]PBR28 imaging detected increased neuroinflammation in CLN7 animals. **(A)** [^11^C]PBR28 SUV imaging of WT and CLN7 animals overlaid on ONPRC18 T_2_w MRI template. **(B)** Histogram displaying [^11^C]PBR28 SUV for WT and CLN7 animals. \**P*<0.05, \*\**P*<0.01. **(C)** Scatterplots of CLN7 animals’ [^11^C]PBR28 SUV vs age with solid lines of best fit and 95% confidence intervals in dashed lines. Abbreviations: FrontC = Frontal Cortex, MotorC = Motor Cortex, TempC = Temporal Cortex, Par-OccC = Parieto-Occipital Cortex, Thal = Thalamus, Caud = Caudate Nucleus, Put = Putamen, Cereb = Cerebellum.

### Biofluid analysis identified markers of neurodegeneration

Finally, a CSF-based biofluid analysis was performed in a subset of animals across a range of ages (see Supplementary Table 4) to compare the profile of NfL, total Tau, GFAP, and UCLH1 between CLN7 and WT animals (Fig. 4A). Statistical analysis with Mann Whitney U test revealed significantly higher levels in CLN7 versus WT animals for NfL (U=0, *P*=0.0007), total Tau (U=6, *P*=0.02), and GFAP (U=8, *P*=0.0426) but not UCHL1 (U=10, *P*=0.0813). For all CLN7 and WT animals, correlations only indicated significant positive associations between NfL and age (CLN7: r=0.90, *P*=0.014; WT: r=0.72, *P*=0.046); and simple linear regression revealed significant differences in the rate of NfL increase with age between the groups (F(1,10)=22.44, *P*=0.0008; WT slope: 11.94, CLN7 slope: 187.0) (Fig. 4B).

**Figure 4.**
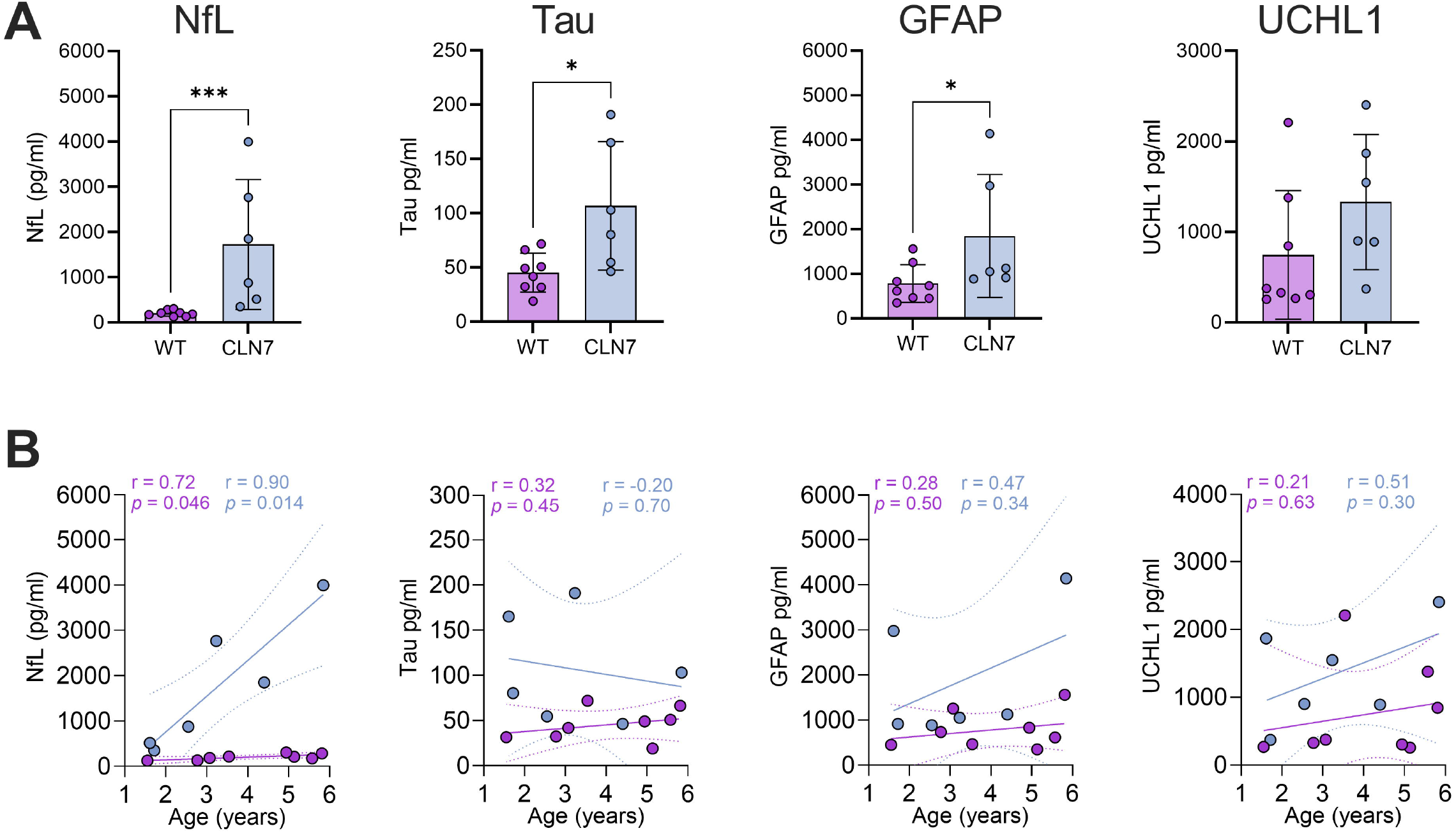
CLN7 animals presented higher levels of neuropathological markers in CSF. **(A)** Histogram of CSF protein analytes measured by ELISA. \**P*<0.05, \*\*\**P*<0.001. (B) Scatterplots of CSF protein analyte levels plotted against age with separate linear regressions for WT and CLN7 animals. Lines of best fit are plotted in solid with 95% confidence intervals as dashed lines. Abbreviations: NfL = Neurofilament light, Tau = Tau Protein, GFAP = Glial Fibrillary Acidic Protein, UCHL1 = Ubiquitin Carboxy-terminal Hydrolase L1.

## Discussion

Clinical understanding of prodromal CLN7 disease remains extremely limited. Although it is a monogenic disorder, routine newborn screening and early genetic testing for CLN7 mutations are not common practice, and patients are rarely diagnosed before the onset of neurological symptoms.^16^ Consequently, compared to other variants, little is known about the sequence of brain and molecular changes that precede overt clinical manifestations.^26,27^ As has been the case for CLN2 disease, defining these early events is essential for establishing sensitive biomarkers and for identifying a therapeutic window during which interventions may prevent or delay irreversible neurodegeneration, and for tracking efficacy of interventions during clinical trials.^28^

This study provides the first multimodal characterization of prodromal and early-stage CLN7 disease in any large animal model. Using structural T_2_w MRI, [^18^F]FDG-PET, [^11^C]PBR28-PET, and CSF biomarkers, we showed that many neurodegenerative and neuroinflammatory changes emerge prior to clinical manifestations. While we previously characterized the gross cerebral and cerebellar atrophy in this spontaneous macaque model,^23^ the analyses reported here reveal a more granular, region-dependent pattern of vulnerability. Several cortical and subcortical regions exhibited early decline during the pre-symptomatic, <2.5 years, and incipient stages, 2.5-3.5 years (e.g. Thalamus, and regions of Occipital, Insula, and Motor cortices), whereas most regions showed measurable atrophy only after 3.5y, coincident with clinical deterioration. While the progressive and regionally selective volume loss observed in CLN7 macaques suggests that neurodegeneration begins subtly during the prodromal phase and accelerates with overt clinical progression, structural MRI alone leaves unanswered questions about whether these anatomical changes reflect metabolic dysfunction, inflammatory processes, or a combination thereof. PET imaging therefore provided complementary information about the functional and inflammatory state of affected tissue.

[^18^F]FDG PET revealed genotype differences in cerebral glucose metabolism that emerged early in regions without significant atrophy (e.g. Frontal and Motor cortical regions, Figure 2B), which expanded to other regions at older ages. Correlational analyses aid in interpreting these genotype differences, where WT animals showed a consistent pattern of positive correlations between age and SUVglc in most of the examined regions that were absent in CLN7 animals. These findings suggest that altered cerebral glucose metabolism during brain development may be an early hallmark of CLN7 disease, preceding, or potentially occurring independently from, gross tissue loss. In contrast, correlation analyses of [^11^C]PBR28 PET data demonstrated widespread age-related increases in neuroinflammation across many of the same regions showing metabolic decline (Figure 3C), and in some cases in regions that remained structurally preserved (e.g. Putamen). These findings suggest that neuroinflammation is a secondary feature of CLN7 pathology, potentially contributing to metabolic impairment and atrophy. Taken together, these data highlight the value of PET imaging for identifying presymptomatic disease involvement and for detecting treatment effects in regions not yet compromised by gross atrophy. In parallel, CSF biomarkers provided convergent evidence of early neurodegenerative pathology. NfL, Tau, and GFAP were elevated in CLN7 animals, and age-associated increases in NfL suggest a cumulative pattern of axonal injury. These findings reinforce the potential of CSF NfL as a sensitive indicator of disease progression.

The sample size represents an unavoidable limitation of this study. *CLN7*^*-/-*^ macaques arise from intentional heterozygote breeding within the ONPRC Japanese macaque colony, but because the mutation is autosomal recessive, only a small number of homozygous *CLN7*^*-/-*^ mutants were available for study. Ethical and welfare considerations further constrain colony expansion. Consequently, only a limited number of subjects could be enrolled, leading to uneven age distributions and our analysis strategy of grouping data into broader age bins. These limitations should be considered when interpreting the results. However, while this constrains statistical power, the reproducible direction and magnitude of effects across modalities (MRI, PET, and CSF) provide strong internal validation of the observed disease trajectories. Nonetheless, future studies with multimodal longitudinal monitoring extending into later symptomatic stages and integrated with behavioral outcomes, as well as complementary post-mortem neuropathological evaluation of prodromal stages, will be valuable to further refine the timeline of disease progression.

The establishment of prodromal biomarkers in a nonhuman primate CLN7^−/−^ model addresses a major gap in the field. In affected children, CLN7 disease is rarely diagnosed before the onset of symptoms, and most available studies describe late-stage features.^16,29^ These clinical observations leave the earliest pathological events unresolved. This naturally occurring macaque model enables assessment of important parameters early in disease progression, providing here the first opportunity to define how structural, metabolic, inflammatory, and neurodegenerative alterations unfold in a brain that closely parallels human anatomy and developmental timing. This alignment offers a uniquely translational platform for evaluating early diagnostic and therapeutic targets.

In conclusion, our study defines, for the first time, a multimodal biomarker signature of CLN7 disease in a nonhuman primate model. Through integrated imaging and CSF analyses, we delineate a cascade of neuroinflammation, metabolic disruptions, and regionally variable degeneration. These findings bridge mechanistic insight and translational relevance, establishing a foundation for early diagnosis, refined disease staging. By mapping when and where neuroinflammation, hypometabolism and atrophy occur, the results from this study inform the design of preventive or disease-modifying interventions in CLN7 disease.

## Data Availability

The data that support the findings of this study are available from the corresponding author upon reasonable request.

## Acknowledgements

We would like to express our gratitude to the ONPRC Division of Comparative Medicine for the excellent care provided to the animals in this study, along with special acknowledgement of the efforts of John Templon, Lauren Martin, Anne Lewis, and Kristin Brandon. We would also like to acknowledge the contributions made by the OHSU Center for Radiochemisty Research on the [^11^C]PBR28 production. The computational resources and services used in this work for the processing of the MRI data were provided by the HPC core facility CalcUA of the University of Antwerp, the VSC (Flemish Supercomputer Center), funded by the Hercules Foundation and the Flemish Government department EWI.

## Funding

National Institutes of Health grants:

K01 AG078407

P51 OD011092

R24 NS104161

MD is supported by the Research Foundation Flanders (FWO, 1156626N).

## Supplementary material

Supplementary material is available at *Brain* online.

