## Supplementary Materials for "Prodromal pathogenesis of CLN7 Batten Disease revealed by multimodal biomarkers in macaques"

### Supplementary Methods

#### Magnetic Resonance Imaging

##### Acquisition

MRI acquisitions were performed at the ONPRC using a Siemens Prisma whole-body 3 Tesla (T) MRI instrument (Erlangen, Germany) with a 16-channel paediatric head RF coil. Animals were initially anesthetized with Ketamine HCl (10-15 mg/kg i.m.), intubated, and then maintained on 1-2% isoflurane gas vaporized in 100% oxygen for the duration of the scan. NHPs were placed in a head-first supine position for the MRI while their heads were immobilized using foam padding to avoid movement artifacts. Heart rate, blood pressure, temperature, and blood oxygenation were monitored throughout the scan by veterinary technicians. 3D $T_2$-weighted ($T_2$w) imaging sequences were acquired with 0.5 mm isotropic voxels (TE/TR = 385/3200 ms, flip angle = 120°, matrix size = 320 × 320, number of slices = 224). Three $T_2$w images were acquired in each MRI session. The total acquisition time of this scan was 29 min 42 s.

##### Processing

$T_2$w images were co-registered and averaged for improved signal quality. Averaged images were registered to the ONPRC18 Whole head template[1], then a brain mask was back-transformed into individual space for skull-stripping and extraction of brain images. To facilitate skull stripping of young animals, a manual brain mask was drawn on one animal under the age of 2y. For all other animals under the age of 2y, averaged images were registered to this scan, and the age-appropriate mask back transformed to each young animal's native space to perform skull-stripping using nonlinear warps. All images were checked for adequate alignment of the transformed whole brain mask to the individual space prior to skull-stripping, and all masks were deemed to be accurate by a blinded experimenter with no manual correction required. An intensity bias correction was then performed on the obtained images ('N4BiasFieldCorrection' tool in ANTs). This specific correction compensates for low-frequency intensity non-uniformity with respect to the coil configuration[2].

### Analysis

To assess macroscopic brain structural alterations, the previously generated macaque brain ONPRC18 T2w template[1] was aligned with nonlinear warps and then the gray matter labelmap was back-transformed to individual subject space (ANTs). No manual adjustments were made to the labelmaps once transformed to individual space. The voxel count of Regions of Interest (ROIs) defined in the ONPRC18 atlas[1], namely DLPFC, dorsolateral prefrontal cortex; VLPFC, ventrolateral prefrontal cortex; OPFC, orbitofrontal cortex; VMPFC, ventromedial prefrontal cortex; DMPFC, dorsomedial prefrontal cortex; ACC, anterior cingulate cortex; DPMC, dorsal premotor cortex; VPMC, ventral premotor cortex; SMC, supplemental motor cortex; MC, primary motor cortex; STC, superior temporal cortex; ITC, inferior temporal cortex; RC, rhinal cortex; IC, insular cortex; SSC, somatosensory cortex; PC, parietal cortex; PCC, posterior cingulate cortex; OCC, occipital cortex; CD, caudate; PUT, putamen; LatTH, lateral thalamus; MdTH, medial thalamus; HIPP, hippocampus; AMY, amygdala; SN, substantia nigra; GPI, internal globus pallidus; GPE, external globus pallidus; and CBM, cerebellum were extracted, bilateral ROIs were combined, and the resulting volume in $mm^3$ used for statistical analysis.

### Positron Emission Tomography (PET) imaging

PET/CT (Computed Tomography) imaging was performed using a GE Discovery MI 710 PET-CT imaging system. Animal preparation was performed as previously reported[3,4]. Animals were initially anesthetized with Ketamine HCl (10-15 mg/kg i.m.), intubated, and then maintained on 1-2% isoflurane gas vaporized in 100% oxygen for the duration of the scan. A saphenous intravenous (i.v.) catheter was placed for ligand administration, arterial line for blood collection, and a cephalic i.v. catheter was established for fluid support. Animals were positioned on the scanner-bed in a head-first prone orientation and immobilized using a PET/CT compatible stereotaxic head frame (Crist Instruments). Heart rate, blood pressure, temperature, and blood oxygenation were monitored throughout the scan by veterinary technicians. For the [$^{18}$F]FDG scan, animals were fasted for approximately 18h prior to the scan. Prior to each PET, an 8-sec CT was collected using 100kV 50mA, for co-registration, attenuation, and scatter correction. Dynamic PET data were collected in list-mode format and reconstructed using the GE software (Q.clear technology). Normalization, scatter, dead time, and CT-based attenuation corrections were applied. PET image frames were reconstructed on a 120x100x52 grid with 1.823x1.823x2.780 $mm^3$ voxels.

**Ligands Radiosynthesis**

To investigate cerebral glucose uptake, [$^{18}$F]FDG PET imaging was performed as previously reported[3]. [$^{18}$F]FDG was obtained commercially from Cardinal Health (Seattle, WA) and produced using a GE PETtrace 880.

To assess neuroinflammation, [$^{11}$C]PBR28 PET imaging was synthesized by the OHSU Center for Radiochemistry Research (CRR). The general chemical scheme for the radiochemical synthesis of [$^{11}$C]PBR28 consists of two parts, the production of $^{11}CH_3I$ (carbon-11 methyliodide) and then the reaction of $^{11}CH_3I$ with the PBR28 precursor to make [$^{11}$C]PBR28. $^{11}CO_2$ was produced *via* $^{14}$N[p, α]$^{11}$C nuclear reaction by bombarding nitrogen gas containing 1% oxygen gas with protons in a GE PETrace 880 16.5 MeV cyclotron. The $^{11}CO_2$ was then passed through a line to an automated GE Healthcare methyliodide production system (TRACERlab FX2 MEI) within a Hot cell. Concisely, the $^{11}CO_2$ was trapped on molecular sieve to isolate the $^{11}CO_2$. Then the gas was released and passed over a nickel catalyst with hydrogen gas at 350°C to make $^{11}CH_4$. The $^{11}CH_4$ was trapped on Carboxen polymer sieve 60-80 mesh, which selectively trapped only the $^{11}CH_4$ but not any remaining $^{11}CO_2$. After trapping, the Carboxen trap was heated and passed in a stream of helium gas to a tube containing iodine at 740°C. Valves were switched so that the gas was recirculated through the iodine oven several times until the maximum of radioactive $^{11}CH_3I$ was achieved. The $^{11}CH_3I$ was trapped on Porapak Q to isolate the $^{11}CH_3I$ then heated and bubbled into a reaction vessel. Next, a reaction vessel containing the PBR28 precursor in dimethylformamide, $^{11}CH_3I$ and 7N NaOH was heated for 5 minutes at 60 °C. The reaction mixture was subsequently purified using preparative high-performance liquid chromatography (HPLC). An isocratic mobile phase, consisting of sterile water for injection and ethanol (57:43, v/v) was used to elute the purified [$^{11}$C]PBR28 from a semi-preparative C18 HPLC column. The [$^{11}$C]PBR28 fraction was collected through a sterilizing filter into a vented sterile vial containing 10mL of 0.9% NaCl saline for injection, USP (preservative free) and 50 $\mu$L sodium phosphates for injection, USP.

#### Blood analysis and plasma correction for [$^{11}$C]PBR28

Characterization of [$^{11}$C]PBR28 blood profile was performed based on serial arterial blood samples collected in parallel to the PET acquisition to calculate arterial input functions and correct for the presence of plasma radiometabolites using a solid-phase extraction method[5]. Briefly, a total of 10 blood samples (1 mL each) at 0.25, 0.5, 0.75, 1, 6, 10, 20, 40, 60, and 80 minutes post injection (p.i.) were collected in 3 mL heparinized syringes. At each timepoint, radioactivity was measured in 200 $\mu$L of whole blood and 100 $\mu$L of plasma in a cross-calibrated gamma counter (Wizard2, PerkinElmer), and used to calculate the plasma to whole blood ratio.

An additional 1.0 mL of arterial blood was collected at 6, 10, 20, 40, 60, and 80 minutes p.i. to measure the parent fraction during the scan and correct the plasma input function for radiometabolites. Following separation via centrifuge (3000xrcf for 5 min), 300 $\mu$L of plasma was pushed through a conditioned SPE cartridge together with 1 mL of micellar eluent, and then 2.0mL d.i. $H_20$. SPE cartridge and flowthrough were measured separately in the gamma counter (Wizard2, PerkinElmer) for quantification of the radiometabolite and parent fractions. The ratio of radioactivity measured in the SPE cartridge, and the eluent determined the percent contribution of the parent ligand to the total radioactivity signal at each sampling time. Using PMOD 3.6 software (Pmod Technologies, Zurich, Switzerland), individual metabolite-corrected [$^{11}$C]PBR28 plasma arterial input function for kinetic modelling of the PET data were obtained by correcting individual input functions by parent fraction values fitted with a sigmoid curve as well as for the plasma-to-whole blood ratios (Supplementary Fig. 1). Due to a methodological issues during blood sampling caused by clogged catheters, in 2 scans it was not possible to collect all of the blood timepoints.

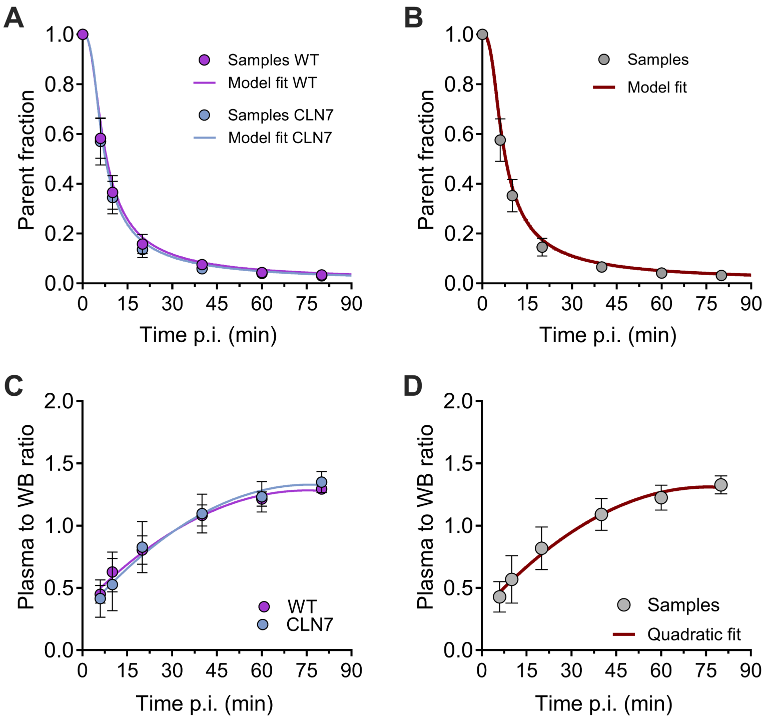


**Supplementary Figure 1. [$^{11}$C]PBR28 Blood kinetic profiles.** (A) Fraction of unmetabolized [$^{11}$C]PBR28 ligand plotted against time after ligand administration with WT and CLN7$^{-/-}$ data plotted separately along with separate sigmoidal models. (B) WT and CLN7$^{-/-}$ parent fraction data plotted together. (C) Ratio of radioactivity in plasma to whole blood plotted against time after ligand administration for WT and CLN7$^{-/-}$ animals along with separate quadratic models. (D) WT and CLN7$^{-/-}$ plasma to whole blood ratio data plotted together.

### Scan Acquisition

[$^{18}$F]FDG PET scan was performed as previously described[3]. Briefly, before the start of the scan, fasting blood glucose levels were measured via a blood sample. Thirty-minutes prior to the onset of the scan, a bolus i.v. injection of ligand was performed manually over a 30s interval. The average injected dose of [$^{18}$F]FDG across scans was 132.6±5.6 MBq. Detailed information on the animal and dosing parameters is provided in the Supplementary Table 2. No significant difference in the radioactive dose administered between groups was observed. [$^{18}$F]FDG PET scans were collected over 50 min reconstructed into 10 frames of 300s each (10x300s).

Upon start of the dynamic [$^{11}$C]PBR28 PET scan acquisition, a bolus of radioligand (190.5 ± 122.46 MBq; injected mass of 0.01510 ± 0.00914 $\mu$g/kg) was injected manually over a 30s interval immediately after the start of the 90 min dynamic PET scan. Detailed information on the animal and dosing parameters is provided in the Supplementary Table 3. No significant difference in the radioactive dose administered between groups was observed. Dynamic [$^{11}$C]PBR28 PET data were reconstructed into 52 frames of increasing length (16×15s, 6×30s, 8×60s, 8×120s, 4×180s, and 10×300s).

### Image processing and analysis

Analysis and processing of PET data were performed using PMOD 3.6 software (Pmod Technologies). Spatial normalization of PET images to the ONPRC18 MRI template[1] was performed using individual $T_2$w MR images using the previously described pipeline[3] for both radioligands. Due to the limited spatial resolution of PET images, Regions of Interest (ROIs) delineated via the ONPRC18 simplified labelmap[1] were used to extract regional time-activity curves (TACs) (Supplementary Figs. 2 and 3).

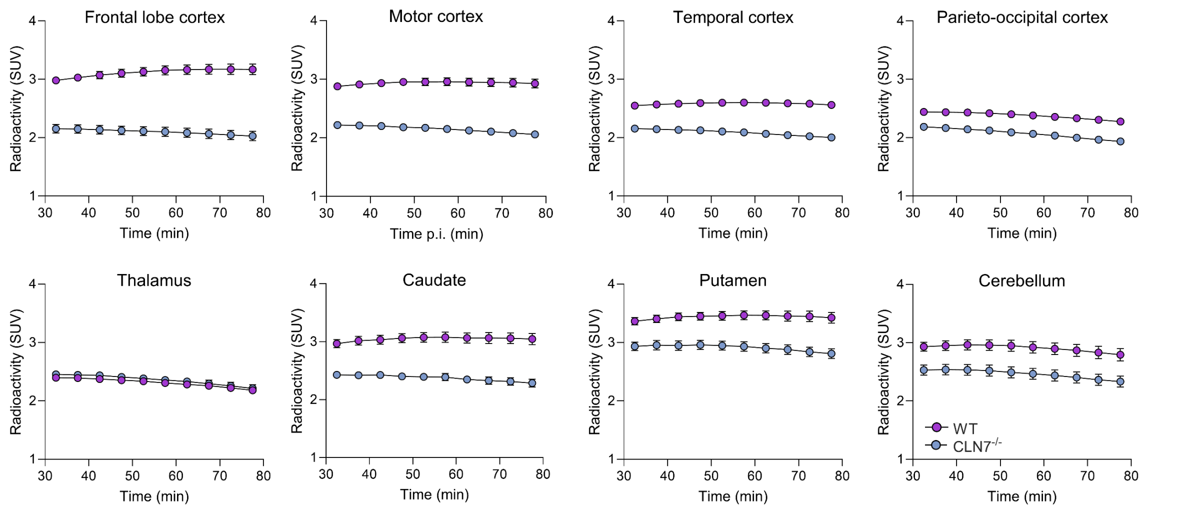


**Supplementary Figure 2. [$^{18}$F]FDG SUV Time Activity Curves.** WT and CLN7$^{-/-}$ SUVs plotted against time after ligand administration for 8 brain regions.

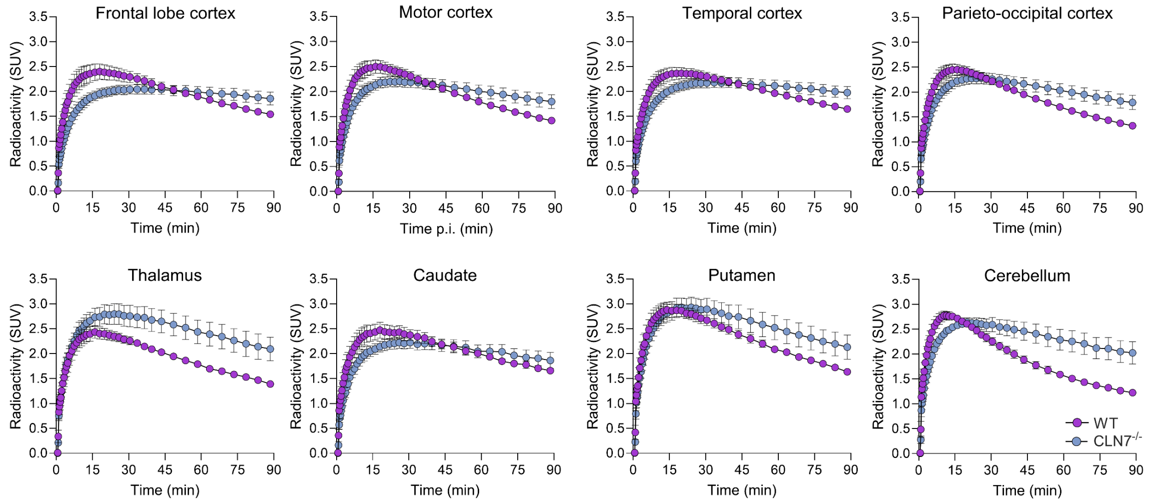


**Supplementary Figure 3. [11C] SUV Time Activity Curves.** WT and CLN7$^{-/-}$ SUVs plotted against time after ligand administration for 8 brain regions.

[$^{18}$F]FDG brain uptake was measured in multiple brain regions by multiplying the SUV (Standardized Uptake Value) with fasted blood glucose levels to obtain the glucose-corrected SUV values ($SUV_{glc}$). SUV values were measured based on the scan interval 60-80 min post-injection of the radioligand. $SUV_{glc}$ [$^{18}$F]FDG PET images were generated in PMOD, represented as group averages, and overlaid onto the ONPRC18 MRI brain template for anatomical reference.

For quantification of [$^{11}$C]PBR28 PET imaging, we explored which tissue compartmental model (TCM) could describe the tracer kinetics and provide an estimate of the volume of distribution ($V_T$) using metabolite-corrected [$^{11}$C]PBR28 plasma arterial input functions. Due to the presence of a known vascular component for [$^{11}$C]PBR28[6], several models including (ir)reversible vascular binding exist. However, after exploration, 2TCM provided the best visual fit in this non-human primate model. In few instances, the model showed large (>25%) standard error estimates of the $V_T$, therefore those values were excluded from the analysis due to low accuracy. As alternative to the 2TCM, this model was subsequently compared to the Logan graphical method[7]. For 2TCM, the blood volume fraction ($V_B$) was fixed at 4%[4], whereas the linear phase (t*) for Logan graphical method was determined from the curve fitting based on 10% maximal error with t* ranging of 34.6 ± 16 min. A strong agreement between $V_T$ estimates based on 2TCM and Logan graphical method was observed (Supplementary Fig. 4). For 2 scans, blood data were not complete and/or available due to clogged arterial catheters, therefore kinetic modelling was not performed. Additionally, $V_T$ estimates displaying a standard error above 30% were excluded from analysis. As an alternative, standardized uptake value (SUV) based on the scan interval 70-90 min post-injection of the radioligand was used for the final analysis given the good agreement with the Logan $V_T$ estimates (Supplementary Fig. 5). Accordingly, SUV [$^{11}$C]PBR28 PET images were generated in PMOD, represented as group averages, and overlaid onto the ONPRC18 MRI brain template for anatomical reference.

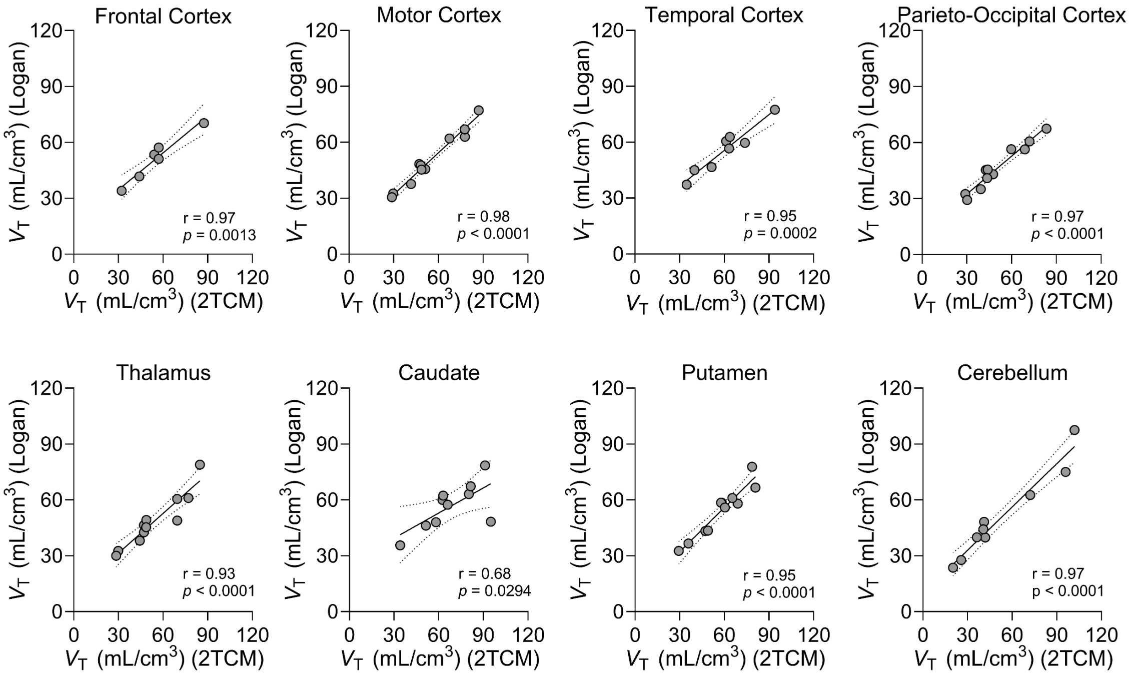


**Supplementary Figure 4. Correlations of 2TCM vs Logan for [$^{11}$C]PBR28.** $V_T$ values calculated from Logan plot displayed on Y-axis with $V_T$ values calculated using a 2TCM plotted on X-axis along with linear regressions as a solid line and confidence interval in dotted line for 8 brain regions.

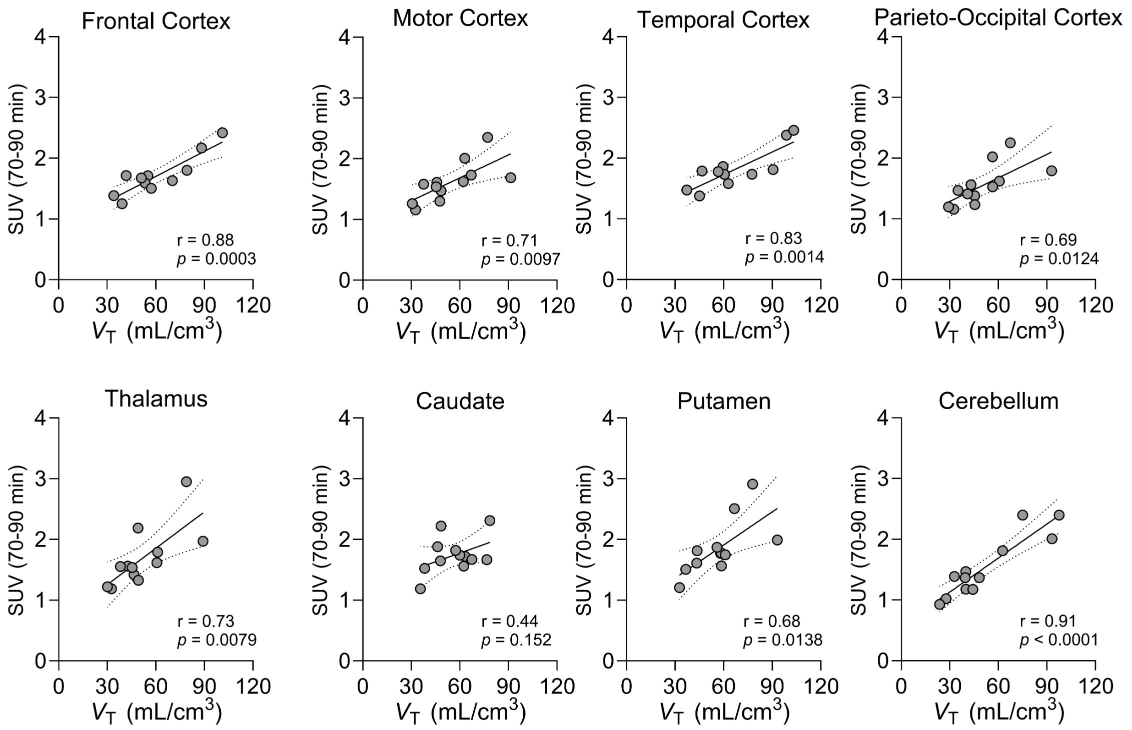


**Supplementary Figure 5. Correlations of $V_T$ vs SUV for [$^{11}$C]PBR28.** SUVs plotted against $V_T$ values calculated from Logan plot along with linear regressions as a solid line and confidence interval in dotted line for 8 brain regions.

#### Cerebrospinal fluid (CSF) biomarkers

In a subset of animals (CLN7$^{-/-}$ *n*=6; WT *n*=8), 1 mL of CSF was collected under isoflurane anaesthesia by cisterna magna aspiration. Samples were immediately frozen on dry ice and stored at -80C before shipping to Quanterix (Massachusetts, USA) where single protein molecule digital ELISA immunoassays using the using the SIMOA neurology 4-plex-A biomarker panel (digital immunoassay) were conducted to measure neurofilament light (Nf-L), total tau (Tau), glial fibrillary acidic protein (GFAP), and ubiquitin carboxyl-terminal hydrolase L1 (UCHL1).

#### Statistical analysis

A two-way ANOVA with Holm-Šidák's multiple comparison test was applied to compare all MRI and PET metrics between experimental groups in each brain structure, a Mann-Whitney test was used to compare analytes between genotype groups in CSF samples. For ANOVAs comparing across age-bins, age bins were defined by grouping animals <2.5 years, 2.5-3.5 years, >3.5 years (MRI); <2.5 years, >2.5 years (FDG). Pearson's correlation tests were used to compute all correlations (MRI, PET, CSF) for each group separately, and simple linear regressions to plot slopes. Statistical analyses were performed using GraphPad Prism (v10) and R (2024.09.0). Data are represented as mean ± standard deviation (SD). All tests were two-tailed and significance was set at $P<0.05$.

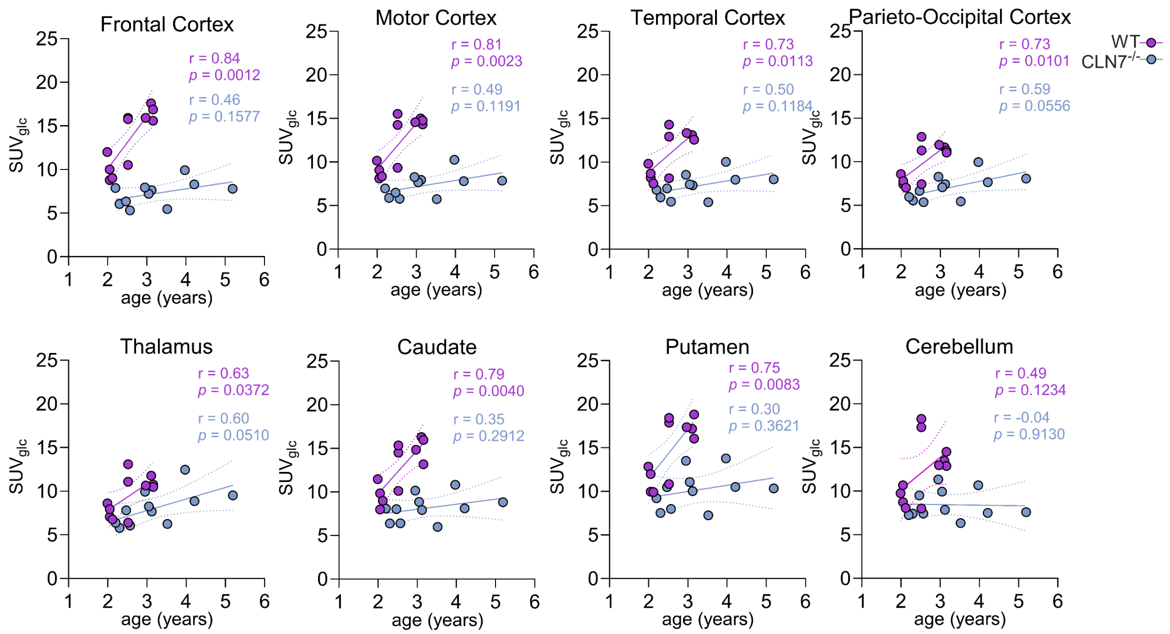


**Supplementary Figure 6. Correlations of [$^{18}$F]FDG SUV$_{glc}$ vs Age.** WT and CLN7$^{-/-}$ [$^{18}$F]FDG SUV$_{glc}$ plotted against age at scan with separate linear regressions as a solid line and confidence intervals as dotted lines for 8 brain regions.

**Supplementary Table 1. [$^{18}$F]FDG Dosing and Demographic Parameters**

| Animal ID / Sex | Age at scan (Years) | Animal Weight (kg) | Activity Administered (MBq) | Fasting Glucose (mmol/l) | Bodyweight per Dose (kg/kBq) |
|---|---|---|---|---|---|
| **CLN7$^{-/-}$** | | | | | |
| M225 / F | 5.19 | 8.35 | 140.008 | 4.33 | 5.96E-05 |
| M226 / M | 3.12 | 6.1 | 131.683 | 3.28 | 4.63E-05 |
| | 3.52 | 6.85 | 134.939 | 2.78 | 5.08E-05 |
| | 3.97 | 9.45 | 137.455 | 3.78 | 6.87E-05 |
| | 4.22 | 9.5 | 130.647 | 4.72 | 7.27E-05 |
| M230 / M | 2.30 | 4.95 | 129.056 | 2.89 | 3.84E-05 |
| | 2.58 | 5.6 | 121.693 | 2.83 | 4.60E-05 |
| | 3.05 | 5.8 | 129.944 | 4.11 | 4.46E-05 |
| M231 / M | 2.20 | 5.45 | 125.356 | 3.00 | 4.35E-05 |
| | 2.47 | 5.9 | 142.524 | 3.22 | 4.14E-05 |
| | 2.95 | 5.8 | 131.128 | 4.39 | 4.42E-05 |
| **WT** | | | | | |
| M233 / M | 1.99 | 4.25 | 128.612 | 3.89 | 3.30E-05 |
| | 3.11 | 6.18 | 133.052 | 5.11 | 4.64E-05 |
| M234 / M | 2.05 | 3.95 | 130.351 | 3.56 | 3.03E-05 |
| | 2.52 | 4.7 | 128.797 | 5.56 | 3.65E-05 |
| | 3.16 | 5.8 | 144.929 | 4.39 | 4.00E-05 |
| M235 / F | 2.05 | 3.65 | 126.577 | 3.50 | 2.88E-05 |
| | 2.52 | 4.25 | 129.426 | 4.83 | 3.28E-05 |
| | 3.16 | 5.65 | 139.12 | 4.83 | 4.06E-05 |
| M236 / F | 2.12 | 3.95 | 136.9 | 2.83 | 2.89E-05 |
| | 2.52 | 4.9 | 128.908 | 3.06 | 3.80E-05 |
| | 2.97 | 5.25 | 136.049 | 5.28 | 3.86E-05 |

**Supplementary Table 2. [$^{11}$C]PBR28 Dosing and Demographic Parameters**

| Animal ID / Sex | Age at scan (Years) | Animal Weight (kg) | Activity Administered (MBq) | Bodyweight per Dose (kg/kBq) | Ligand Mass ($\mu$g) | Ligand Mass per bodyweight ($\mu$g/kg) |
|---|---|---|---|---|---|---|
| **CLN7$^{-/-}$** | | | | | | |
| M225 / F | 5.25 | 7.45 | 112.258 | 4.03E-05 | 0.05026 | 0.00675 |
| M226 / M | 3.17 | 6.3 | 162.027 | 4.16E-05 | 0.11593 | 0.01840 |
| | 3.66 | 6.85 | 245.421 | 6.64E-05 | 0.06661 | 0.00972 |
| | 4.34 | 8.7 | 159.988 | 3.89E-05 | 0.06412 | 0.00737 |
| | 3.96 | 9.7 | 232.545 | 2.79E-05 | 0.11940 | 0.01231 |
| M230 / M | 2.71 | 5.65 | 157.768 | 5.44E-05 | 0.08391 | 0.01485 |
| | 3.07 | 5.9 | 68.036 | 4.17E-05 | 0.12519 | 0.02122 |
| M231 / M | 2.61 | 6 | 232.456 | 3.58E-05 | 0.08531 | 0.01422 |
| | 3.04 | 6.25 | 174.181 | 8.67E-05 | 0.10489 | 0.01678 |
| **WT** | | | | | | |
| M233 / M | 2.01 | 4.2 | 285.751 | 1.47E-05 | 0.10181 | 0.02424 |
| | 2.49 | 4.75 | 174.048 | 2.73E-05 | 0.05763 | 0.01213 |
| M234 / M | 2.57 | 4.4 | 235.283 | 1.87E-05 | 0.07470 | 0.01698 |
| M235 / F | 2.53 | 4.25 | 246.235 | 1.73E-05 | 0.08267 | 0.01945 |
| M236 / F | 2.59 | 4.9 | 135.531 | 3.62E-05 | 0.06434 | 0.01313 |
| | 2.97 | 5.5 | 235.664 | 2.33E-05 | 0.10282 | 0.01869 |

**Supplementary Table 3. MRI Demographic Parameters**

| Animal ID / Sex | Age at scan (Years) |
|---|---|
| **$CLN7^{-/-}$** | |
| M225 / F | 5.19 |
| M226 / M | 3.13 |
| | 3.62 |
| | 3.94 |
| | 4.21 |
| M230 / M | 2.25 |
| | 2.55 |
| | 3.13 |
| M231 / M | 2.15 |
| | 2.45 |
| | 3.02 |
| **WT** | |
| M233 / M | 1.94 |
| | 2.49 |
| | 3.1 |
| M236 / F | 2.07 |
| | 2.6 |
| | 2.98 |
| M234 / M | 1.99 |
| | 2.55 |
| | 3.16 |
| M235 / F | 1.99 |
| | 2.55 |
| | 3.16 |
| M228 / M | 1.4 |
| M221 / M | 4.94 |
| M218 / F | 5.13 |
| M229 / M | 3.07 |
| M217 / F | 5.81 |
| M227 / M | 1.4 |
| M224 / M | 3.1 |

**Supplementary Table 4. CSF Sample Collection Parameters**

| CSF Sample Collection Parameters | |
|---|---|
| Animal ID / Sex | Age at collection (Years) |
| **$CLN7^{-/-}$** | |
| M220 / M | 4.4 |
| M226 / M | 2.55 |
| M230 / M | 1.72 |
| M231 / M | 1.61 |
| **WT** | |
| M238 / F | 5.84 |
| M239 / M | 3.23 |
| M221 / M | 4.94 |
| M218 / F | 5.13 |
| M229 / M | 3.07 |
| M217 / F | 5.81 |
| M227 / M | 1.4 |
| M240 / M | 5.57 |
| M241 / F | 3.54 |
| M242 / F | 2.77 |
| M243 / F | 1.55 |

### Supplementary Table 5. MRI Volumetric ANOVAs with Multiple Comparisons

| ROI | Main effect of Genotype | | Main effect of Age Bin | | Interaction | | Post-hoc (CLN7 vs WT) | | |
|---|---|---|---|---|---|---|---|---|---|
| | | | | | | | <2.5y | 2.5y-3.5y | >3.5y |
| | *F (1, 38)* | *p-value* | *F (2,38)* | *p-value* | *F(2,38)* | *p-value* | *p-value* | *p-value* | *p-value* |
| DLPFC | 10.3318 | **0.0027** | 4.7048 | **0.0149** | 4.3398 | **0.0201** | 0.9315 | 0.1381 | **0.0010** |
| VLPFC | 20.1078 | **0.0001** | 5.9550 | **0.0056** | 5.9839 | **0.0055** | 0.7865 | **0.0204** | **0.0001** |
| OPFC | 12.1000 | **0.0013** | 3.8117 | **0.0310** | 3.8281 | **0.0306** | 0.5247 | 0.2765 | **0.0006** |
| VMPFC | 7.2910 | **0.0103** | 0.7321 | 0.4875 | 5.9940 | **0.0055** | 0.6631 | 0.6081 | **0.0004** |
| DMPFC | 3.0688 | 0.0879 | 5.8183 | **0.0062** | 7.8660 | **0.0014** | 0.1381 | 0.9269 | **0.0006** |
| ACC | 14.8839 | **0.0004** | 5.6732 | **0.0070** | 6.5932 | **0.0035** | 0.8227 | 0.0655 | **0.0001** |
| DPMC | 18.6478 | **0.0001** | 1.5154 | 0.2327 | 2.0340 | 0.1448 | **0.0230** | 0.1949 | **0.0010** |
| VPMC | 38.2201 | **<0.0001** | 1.1776 | 0.3190 | 3.9712 | **0.0272** | **0.0471** | **0.0021** | **<0.0001** |
| SMC | 1.6649 | 0.2047 | 17.1995 | **<0.0001** | 6.4054 | **0.0040** | 0.0670 | 0.7935 | **0.0041** |
| MC | 1.4419 | 0.2373 | 0.0719 | 0.9308 | 0.6772 | 0.5141 | 0.3136 | 0.8181 | 0.2416 |
| STC | 16.5762 | **0.0002** | 3.4358 | **0.0425** | 9.8289 | **0.0004** | 0.5014 | 0.0694 | **<0.0001** |
| ITC | 9.9772 | **0.0031** | 1.4357 | 0.2506 | 5.1892 | **0.0102** | 0.8213 | 0.2186 | **0.0005** |
| RC | 5.1519 | **0.0290** | 3.8012 | **0.0313** | 3.0913 | 0.0570 | 0.6125 | 0.2006 | **0.0109** |
| IC | 15.9980 | **0.0003** | 0.5621 | 0.5747 | 4.0701 | **0.0250** | 0.6292 | **0.0458** | **0.0004** |
| SSC | 18.7768 | **0.0001** | 0.6178 | 0.5445 | 3.1265 | 0.0553 | 0.2358 | 0.0570 | **0.0003** |
| PC | 10.0153 | **0.0031** | 6.9787 | **0.0026** | 4.6664 | **0.0154** | 0.6776 | 0.0587 | **0.0015** |
| PCC | 11.7450 | **0.0015** | 1.2396 | 0.3009 | 6.2771 | **0.0044** | 0.6814 | 0.1229 | **0.0002** |
| OCC | 21.6629 | **<0.0001** | 0.7395 | 0.4841 | 3.5608 | **0.0382** | 0.3623 | **0.0076** | **0.0003** |
| CD | 0.3142 | 0.5784 | 0.3196 | 0.7284 | 3.7009 | **0.0340** | 0.1607 | 0.0550 | 0.0862 |
| PUT | 2.4660 | 0.1246 | 1.9292 | 0.1592 | 0.8964 | 0.4165 | 0.4115 | 0.9969 | 0.0989 |
| GPE | 4.0492 | 0.0513 | 0.1112 | 0.8950 | 1.0650 | 0.3548 | 0.7271 | 0.4114 | **0.0485** |
| GPI | 4.6686 | **0.0371** | 0.3681 | 0.6945 | 1.0701 | 0.3531 | 0.8316 | 0.1736 | 0.0619 |
| SN | 1.5050 | 0.2275 | 2.3307 | 0.1110 | 0.5597 | 0.5760 | 0.2563 | 0.1529 | 0.8580 |
| THAL | 61.1888 | **<0.0001** | 2.6120 | 0.0865 | 0.5922 | 0.5581 | **0.0001** | **<0.0001** | **<0.0001** |
| HIPP | 2.1194 | 0.1537 | 4.1891 | **0.0227** | 3.8815 | **0.0292** | 0.6362 | 0.6659 | **0.0077** |
| AMY | 9.4205 | **0.0039** | 0.8328 | 0.4426 | 5.7431 | **0.0066** | 0.6875 | 0.2568 | **0.0004** |
| CBM | 12.7562 | **0.0010** | 0.9493 | 0.3960 | 5.8374 | **0.0062** | 0.8365 | 0.1042 | **0.0002** |

### Supplementary Table 6. MRI Volumetric Correlations

| ROI | CLN7 correlation w/ Age | | WT correlation w/Age | |
|---|---|---|---|---|
| | Pearson r | p-value | Pearson r | p-value |
| DLPFC | -0.8048 | **<0.0001** | 0.0100 | 0.9675 |
| VLPFC | -0.8244 | **<0.0001** | -0.0142 | 0.9541 |
| OPFC | -0.8867 | **<0.0001** | 0.0314 | 0.8986 |
| VMPFC | -0.8123 | **<0.0001** | 0.2852 | 0.2365 |
| DMPFC | -0.8649 | **<0.0001** | 0.0945 | 0.7004 |
| ACC | -0.8600 | **<0.0001** | 0.0907 | 0.7120 |
| DPMC | -0.6371 | **0.0006** | -0.0473 | 0.8474 |
| VPMC | -0.7406 | **<0.0001** | 0.1851 | 0.4480 |
| SMC | -0.9052 | **<0.0001** | -0.2798 | 0.2459 |
| MC | -0.2346 | 0.2589 | 0.0346 | 0.8881 |
| STC | -0.8734 | **<0.0001** | 0.2098 | 0.3885 |
| ITC | -0.7564 | **<0.0001** | 0.2474 | 0.3072 |
| RC | -0.8251 | **<0.0001** | 0.0463 | 0.8507 |
| IC | -0.6156 | **0.0011** | 0.3100 | 0.1965 |
| SSC | -0.6479 | **0.0005** | 0.1801 | 0.4606 |
| PC | -0.8748 | **<0.0001** | -0.1033 | 0.6738 |
| PCC | -0.7764 | **<0.0001** | 0.3109 | 0.1952 |
| OCC | -0.5937 | **0.0018** | 0.2597 | 0.2830 |
| CD | -0.3832 | 0.0587 | 0.4209 | 0.0728 |
| PUT | 0.0075 | 0.9716 | 0.5472 | **0.0153** |
| GPE | -0.1766 | 0.3984 | 0.4405 | 0.0591 |
| GPI | -0.2124 | 0.3081 | 0.3554 | 0.1353 |
| SN | 0.5113 | **0.0090** | 0.2665 | 0.2702 |
| THAL | 0.1086 | 0.6053 | 0.3947 | 0.0945 |
| HIPP | -0.0891 | 0.6719 | 0.6457 | **0.0028** |
| AMY | -0.9047 | **<0.0001** | 0.3361 | 0.1595 |
| CBM | -0.8125 | **<0.0001** | 0.2435 | 0.3152 |
